# Growth differentiation factor-15 regulates oxLDL-induced lipid homeostasis and autophagy in human macrophages

**DOI:** 10.1101/354043

**Authors:** Kathrin Ackermann, Gabriel A. Bonaterra, Ralf Kinscherf, Anja Schwarz

## Abstract

Growth differentiation factor-15 (GDF-15), a divergent and distant member of the transforming growth factor-β superfamily, is suggested as a risk factor for cardiovascular diseases. Thus, we are interested to investigate the influence of GDF-15 in lipid homeostasis and autophagy in macrophages (MΦ) during foam cell formation. Our investigations represent the impairment of GDF-15 on modulators of autophagy and lipid homeostasis in PMA-differentiated human THP-1 MΦ. In this context, *in vitro* resulted GDF-15 silencing in a reduction of lipid accumulation, whereas the addition of recombinant (r)GDF-15 increased the lipid accumulation in human MΦ independent of oxidized (ox)LDL. Additionally, GDF-15 affected the expression of autophagy-relevant proteins (p62, Atg5 and Atg12/Atg5 protein complex) and the p62 accumulation in THP-1 MΦ. Hence, our data suggest that GDF-15 is involved in the regulation of the lipid homoeostasis of human MΦ by regulating autophagic processes.

## Introduction

Atherosclerosis, a chronic inflammation in the arterial wall, is accompanied by accumulation of lipids, inflammatory cells and foam cell formation. Studies have shown, that elevated circulating levels of GDF-15 reflect endothelial activation and vascular inflammation, pathways regulating development and progression of atherosclerosis (Schober ***et al***, 2001). In general, increased serum cholesterol levels and especially low-density lipoprotein (LDL) cholesterol in the form of oxidized low-density lipoproteins (oxLDL), as well as macrophages (MΦ) play a key role in atherosclerotic processes, because MΦ internalize oxLDL via scavenger receptors and develop as foam cells. In this context, growth differentiation factor-15 (GDF-15) is a divergent and distant member of the TGF-β superfamily, identical to MΦ-inhibitory cytokine-1 (MIC-1) (Bootcov ***et al***, 1997; Fairlie ***et al***, 1999). A wide range of cell types express GDF-15, which acts as a pleiotropic cytokine and is involved in the response of cells after oxidative stress and cellular injury. GDF-15 is widely distributed in adult tissues, expressed in epithelial cells and MΦ (Böttner ***et al***, 1999b, Unsicker ***et al***, 2013). GDF-15 expression is upregulated in mononuclear cells and/or MΦ by a variety of stimuli including interleukin (IL)-1β, IL-2, tumor necrosis factor alpha (TNFα), phorbol myristate acetate (PMA), retinoic acid (RA), ceramide, oxLDL, hydrogen peroxide or MΦ-colony-stimulating factor (M-CSF). Thus, GDF-15 seems to be associated with MΦ activation (Bootcov ***et al***, 1997; Fairlie ***et al***, 2001; Schober ***et al***, 2001; Schlittenhardt ***et al***, 2004). Though, increased GDF-15 protein levels are also associated with disease states such as tissue hypoxia, inflammation, acute injury and oxidative stress (Unsicker ***et al***, 2013). Therefore, it is noteworthy to mention that recent and older studies have shown that healthy women with higher serum GDF-15 levels are at increased risk of having cardiovascular events (Brown ***et al***, 2002; Lin ***et al***, 2014). However, the role of GDF-15 during atherosclerotic processes is controversially discussed. On the one hand, GDF-15 deficiency has been shown to inhibit atherosclerosis progression in ApoE^−/−^ mice after 20 weeks of a high-fat diet (Bonaterra ***et al***, 2012). On the other hand, transgenic overexpression of GDF-15 has been demonstrated to play a protective role in advanced stages of atherosclerosis in the ApoE^−/−^ mouse model (after 6 months of a high-fat diet) by observing the limitation or repair of atherosclerotic lesions (Johnen ***et al***, 2012). Moreover, *in vitro* analyses of oxLDL-treated human MΦ showed an increased GDF-15 expression in addition to foam cell formation and apoptosis (Schlittenhardt ***et al***, 2004). In processes of MΦ-derived foam cell formation and cell fate are associated with autophagy (Liu ***et al***, 2016), which in general is an evolutionarily conserved mechanism and is important for numerous physiological and pathological processes (Mizushima ***et al***, 2011). Of note, clinical studies have shown a co-localization of autophagy-related proteins with MΦ in atherosclerotic plaques (Razani ***et al***, 2012; Zhou ***et al***, 2016). Nevertheless, the role of autophagy in development and progression of atherosclerosis seems to be complex. It is fact, that in advanced stages of atherosclerosis, autophagy is impaired (Razani ***et al***, 2012). In this context, animal experiments using mice with MΦ-specific deletion of autophagy gene Atg5 have recently shown that these mice developed plaques with increased apoptosis, as well as oxidative stress and exhibited enhanced plaque necrosis (Liao ***et al***, 2012).

Because the role of GDF-15 during development of atherosclerosis is diversely discussed (Bonaterra ***et al***, 2012, Johnen ***et al***, 2012; Preusch ***et al***, 2013) and a relation between GDF-15, apoptosis, as well as autophagy and atherosclerosis development/progression may be suggested due to histomorphometrical data of atherosclerotic lesions (Bonaterra ***et al***, 2012). We aimed to decipher the role of GDF-15 on formation of foam cells with special respect to autophagy in oxLDL-treated human MΦ.

## Results

### Impairment of oxLDL on GDF-15 expression in human THP-1 MΦ

To investigate the effects of GDF-15 on autophagic processes and lipid accumulation, we added recombinant (r)GDF-15 or transiently silenced GDF-15 expression in PMA-differentiated human THP-1 MΦ (siGDF-15-MΦ) respectively (Fig 1). As negative control, we used a non-silencing siRNA (nsiGDF-15) without homology on any known mammalian gene. rGDF-15 resulted in an increase of cellular GDF-15 from 36.9 ± 6.5 pg/μg protein to 58.2 ± 9.5 pg/μg protein (p=0.09 vs. medium control) using 1.0 μg/ml rGDF-15 or 65.3 ± 10.1 pg/μg protein (p=0.03 vs. medium control) using 1.5 μg/ml rGDF-15 (Fig. 1A). The transient transfection achieved a significant (p<0.03 vs. nsiGDF-15) knock-down of GDF-15, i.e. GDF-15 mRNA level decreased from 100 % to 25 % ± 1.7 % (−75 %; p=0.029) (Fig 1B) and protein level decreased from 44.2 ± 9.0 pg/μg protein to 20.9 ± 1.9 pg/μg protein (−53 %; p=0.026) compared with control (nsiGDF-15) (Fig 1C). Additionally, exposure of nsiGDF-15 MΦ to 50 μg/ml oxLDL for 4h resulted in a 37 % ± 12 % (p<0.001) higher GDF-15 mRNA expression in comparison with medium control (Fig 1B), whereas this significant oxLDL-induced increase in GDF-15 mRNA level was not seen in siGDF-15 MΦ in comparison with siGDF-15 MΦ cultured in medium alone (Fig 1B). However, the exposure (4h) of THP-1 MΦ, siGDF-15- or nsiGDF-15 MΦ to 50 μg/ml oxLDL had no effect on the GDF-15 protein level (Fig 1A, C).

**Figure 1.**
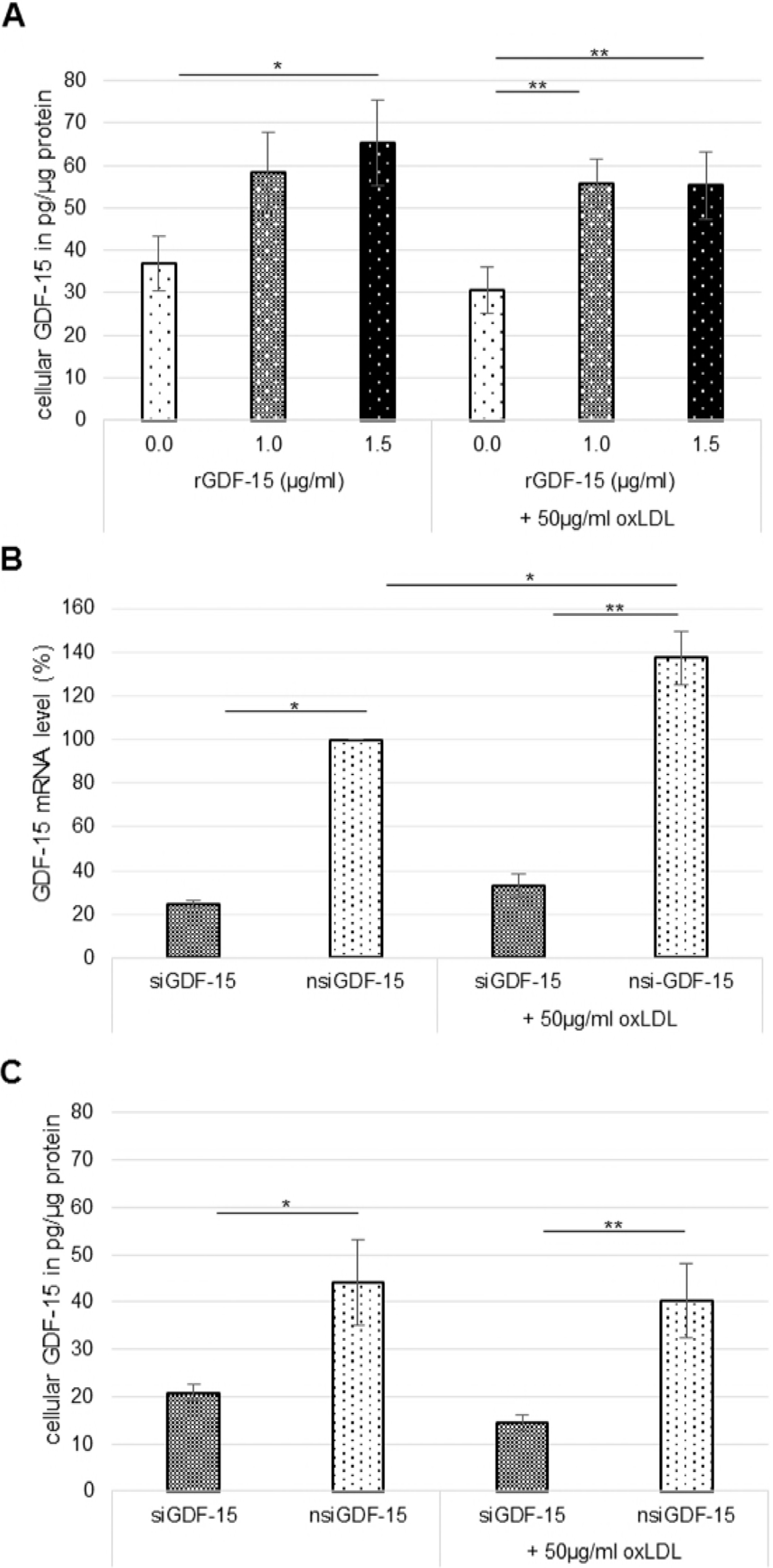
GDF-15 mRNA and protein level in human THP-1 MΦ. THP-1 MΦ were incubated with oxLDL (50 μg/ml), rGDF-15 (1.0 μg/ml, 1.5 μg/ml), oxLDL/rGDF-15 or without (medium control) (A). In addition siGDF-15 or nsiGDF-15 THP-1 MΦ were incubated without (medium control) or with 50 μg/ml oxLDL (B-C). A,C The cellular GDF-15 level level was determined using ELISA [OD_490/655_] and expressed as pg/μg protein (n=5-6 independent experiments). B The GDF-15 mRNA expression levels (in %) were determined using RT-qPCR (n=3 independent experiments). Data information: Bars represent mean ± SEM. *p≤0.05 significance vs. medium control, **p<0.001 significance vs. oxLDL-treated MΦ; (One Way ANOVA; Holm-Sidak method)

### GDF-15 impairs the cellular lipid content in human THP-1 MΦ

Mice MΦ are able to process intracellular lipids via the autophagy-lysosome pathway (Ouimet ***et al***, 2011). For this reason, we analyzed by ORO-staining, whether the intracellular lipid-storage is dependent from GDF-15 in human THP-1 MΦ. After addition of rGDF-15, the cellular lipid content in THP-1 MΦ significantly (p<0.05) increased (1.0 μg/ml rGDF-15:109 % ± 3.9 %; 1.5 μg/ml rGDF-15: 117 % ± 3.7 %) compared to THP-1 MΦ cultured in medium alone (Fig. 2A, C-D). Vice versa, in siGDF-15 MΦ cultured in medium, we found a significant (p=0.02) −19 % decrease of the cellular lipid content (Fig 2B, G-H). Moreover, 50 μg/ml oxLDL (4h) treatment resulted in a significant (p<0.01) 137 % ± 7.1 % increase of the lipid accumulation (Fig 2A, C-F). Additionally, oxLDL treatment led to a significant (p≤0.028) advanced accumulation of lipids to nsiGDF-15 THP-1 MΦ (129 % ± 10.9 %) (Fig 2B, H-J) and siGDF-15 THP-1 MΦ (111 % ± 9.6 %) (Fig 2B, G-I), whereas oxLDL-treated siGDF-15 MΦ thereby showing a significant −18 % (p<0.05) lower lipid accumulation compared to oxLDL-treated nsiGDF-15 MΦ (Fig 2B, I-J). Certainly, THP-1 MΦ co-incubated with rGDF-15/oxLDL did not change the lipid content significantly compared to oxLDL-treated MΦ (Fig 2A, D-F).

**Figure 2.**
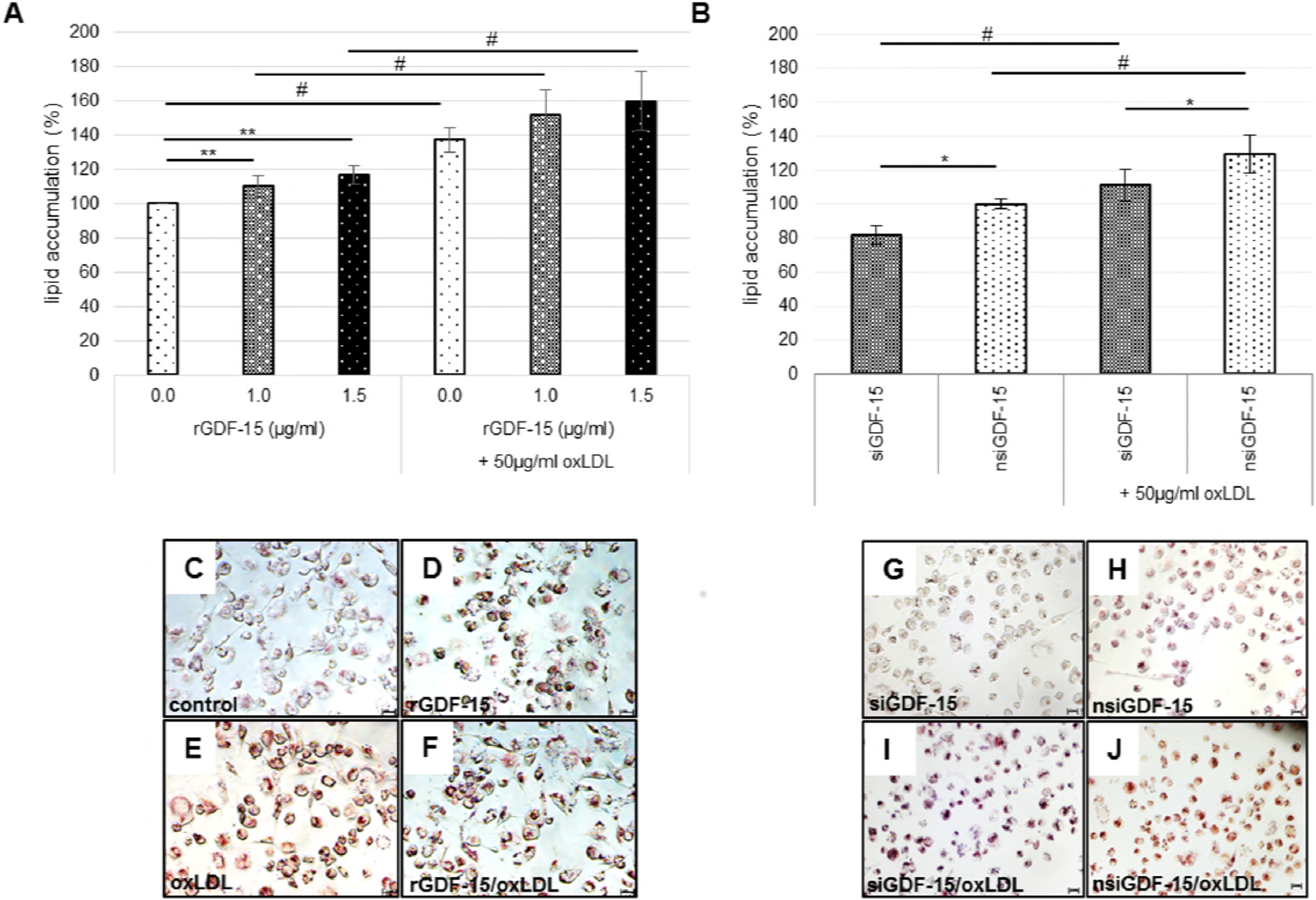
GDF-15 influences the lipid accumulation in THP-1 MΦ. THP-1 MΦ were incubated with oxLDL (50 μg/ml), rGDF-15 (1.0 μg/ml, 1.5 μg/ml), oxLDL/rGDF-15 or without (medium control) (A, C-F). In addition siGDF-15 or nsiGDF-15 THP-1 MΦ were incubated without (medium control) or with 50 μg/ml oxLDL (B, G-J). A-B The ORO-staining of accumulated lipids (%) was quantified and measured after isopropanol elution [OD_510_]. C-J Cells were stained with OilRedO and image by interference light microscopy. Scale bars: C-F: 100 μm; G-J: 20 μm Data information: (A-B) Data presented as mean ± SEM of 6-7 independent experiments. *p≤0.05 significance vs. nsiGDF-15, ^#^p≤0.05 significance vs. without oxLDL; **p≤0.05 significance vs. medium control (Student-Newman-Keuls Method)

### GDF-15 affects the ATG5 protein level and p62 accumulation in human THP-1 MΦ

Autophagic MΦ are found at sides of atherosclerotic lesions, in which the autophagy flux is impaired with lesion progression, indicating a crucial role for autophagy in atherosclerosis (Razani ***et al***, 2012; Zhou ***et al***, 2016). Regarding the link between lipid accumulation and autophagy, we further investigated a potential role of GDF-15 on autophagy. Therefore, we examined the expression of autophagy-relevant proteins/complexes ATG5, ATG12/ATG5 and p62 in THP-1 MΦ dependent from GDF-15 and oxLDL.

Treatment of THP-1 MΦ with oxLDL or rGDF-15 for 4h did not affect the ATG5 and ATG12/ATG5-complex protein (Fig 3A, B). Amazingly the co-incubation of THP-1 MΦ with oxLDL/rGDF-15 (4h) showed a significant (p≤0.04) increase (1.0 μg/ml rGDF-15: 49 %; 1.5 μg/ml rGDF-15: 47 %) of ATG5 protein level (Fig 3A) compared to rGDF-15-treatment alone. In context, ATG12/ATG5 protein complex increased (1.0 μg/ml rGDF-15: 107 %; 1.5 μg/ml rGDF-15: 141 %; p≤0.03) significantly by co-incubation of THP-1 MΦ with oxLDL/rGDF-15 (4h) compared to rGDF-15-treated MΦ (Fig 3B).

Down-regulation of cellular GDF-15 in human THP-1 MΦ (siGDF-15 MΦ) resulted in a significant, oxLDL-independent reduction of ATG5 protein (medium-treated siGDF-15 MΦ by −35 % [p<0.05]; oxLDL-treated siGDF-15 MΦ by −33 % [p=0.03]) compared to nsiGDF-15-MΦ (Fig 3C). ATG12/ATG5-complex was reduced by −15 % (p=0.03) in siGDF-15 MΦ compared to nsiGDF-15 MΦ (100 % ± 2.6 %) (Fig 3D).

**Figure 3.**
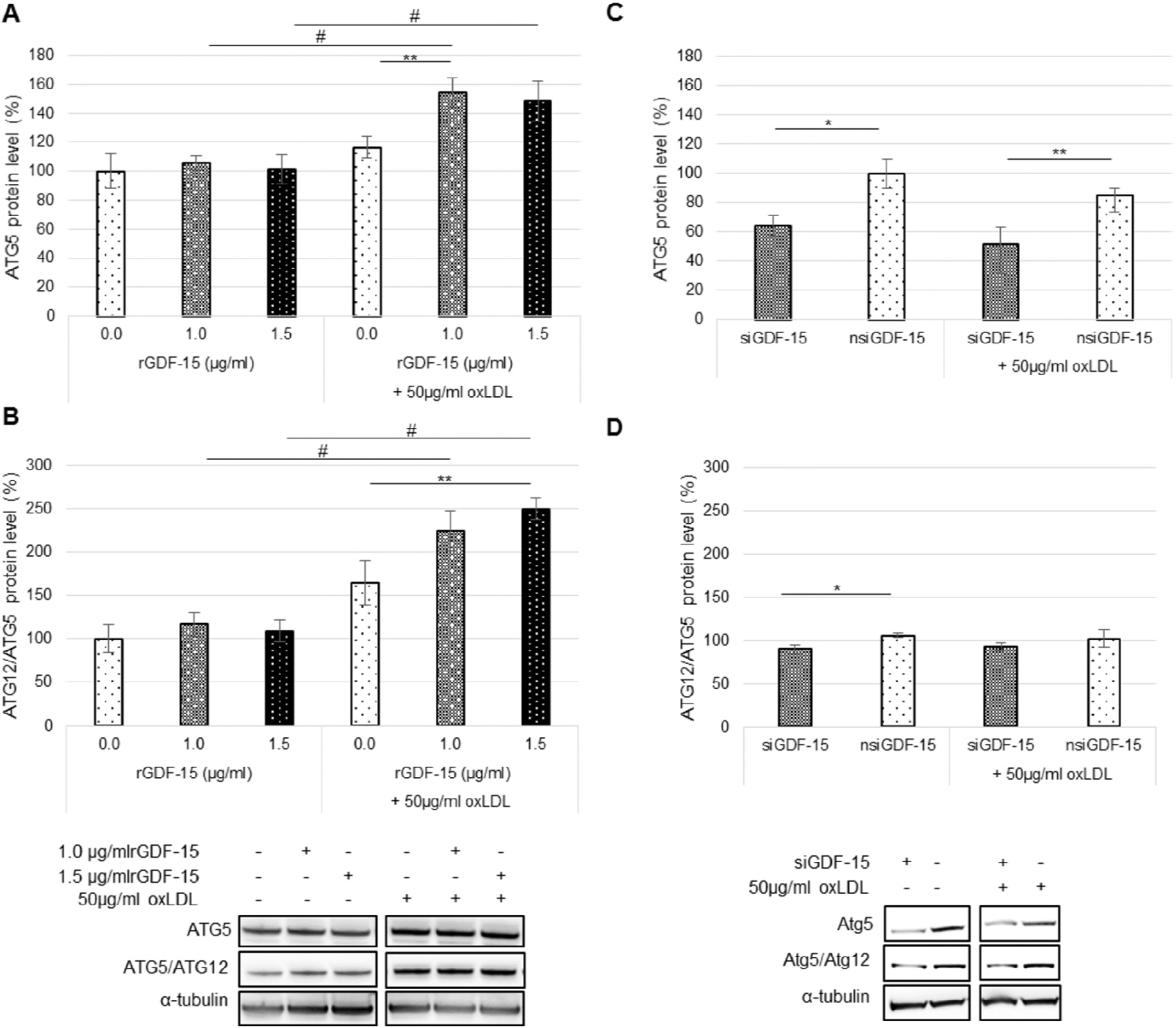
GDF-15 influences the expression of autophagy-relevant ATG5 and ATG12/ATG5-complex in THP-1 MΦ. THP-1 MΦ were treated with oxLDL (50 μg/ml), rGDF-15 (1.0 μg/ml, 1.5 μg/ml), oxLDL/rGDF-15 or without (medium control) (A, B). In addition siGDF-15 or nsiGDF-15 THP-1 MΦ were incubated without (medium control) or with 50 μg/ml oxLDL (C, D). A,C Atg5 and α-tubulin protein levels (%) were determined using western blot analysis and quantified by ImageJ. Expression was normalized against α-tubulin. B,D Atg12/Atg5-complex and α-tubulin protein levels (%) were determined using western blot analysis and quantified by ImageJ. Expression was normalized against α-tubulin. Data information: Bars represent mean ± SEM of 4-6 independent experiments. *p<0.05 significance vs. nsiGDF-15, ^#^p≤0.04 significance vs. without oxLDL; **p≤0.05 significance vs. oxLDL-treated MΦ (One Way ANOVA; Holm-Sidak-Method). *Source data are available online for this figure.*

As a marker of autophagic status, p62-protein increased in oxLDL-treated THP-1 MΦ (THP-1 MΦ: 131 % ± 16.2 % [p=0.065]; nsiGDF-15 THP-1 MΦ: 161 % ± 26.0 % [p=0.03]) (Fig 4A, B). 1.5 μg/ml rGDF-15 co-incubated with oxLDL further amplified the significant increase of p62-protein about 66 % (p<0.05) compared to oxLDL treatment (Fig 4A). In this context, without oxLDL siGDF-15 THP-1 MΦ showed a significant (p<0.05) −33 % reduced p62-protein level compared to nsiGDF-15 THP-1 MΦ (Fig 4B). In addition to our western blot data, we immunocytochemically investigated the intracellular p62 localization in THP-1 MΦ using confocal laser scanning microscopy. We noted that endogenously expressed p62 was present in numerous round bodies (mean size: ≥1.7 μm) in the perinuclear area and a diffuse distribution in cytoplasm of human THP-1 MΦ (Fig 4C-L). For the assessment of the autophagy flux, we analyzed the p62 accumulation in THP-1 MΦ with or without (~control) exposure to oxLDL. Starvation analyses of THP-1 MΦ as a positive control for a suppressed autophagy flux showed an increased p62-accumulation (Fig 4H). 1.5 μg/ml rGDF-15 alone increased (126 % ± 14.4 %; p<0.05) significantly the p62-accumulation in THP-1 MΦ compared to medium control (Fig 4C, D, M). OxLDL-treated THP-1 MΦ verifies the significant 148 % ± 14.8 % increase, whereas 1.5 μg/ml rGDF-15 enhanced the p62 accumulation of 32 % (Fig 4E, F, M). In addition, we found that the p62 accumulation was significantly (p<0.002) reduced by −56 % in siGDF-15 MΦ (Fig 4I, J, N). OxLDL treatment increased the p62 accumulation in nsiGDF-15 and siGDF-15 THP-1 MΦ (nsiGDF-15 MΦ: to 179 % ± 22 %, p=0.01; siGDF-15 MΦ: to 111 % ± 15 %, p=0.013) (Fig 4K, L, N). Linked to the previous data siGDF-15 MΦ treated with 50 μg/ml oxLDL showed a significantly (p<0.001) −64 % reduced p62-protein accumulation compared to nsiGDF-15 MΦ (Fig 4I-L, N).

**Figure 4.**
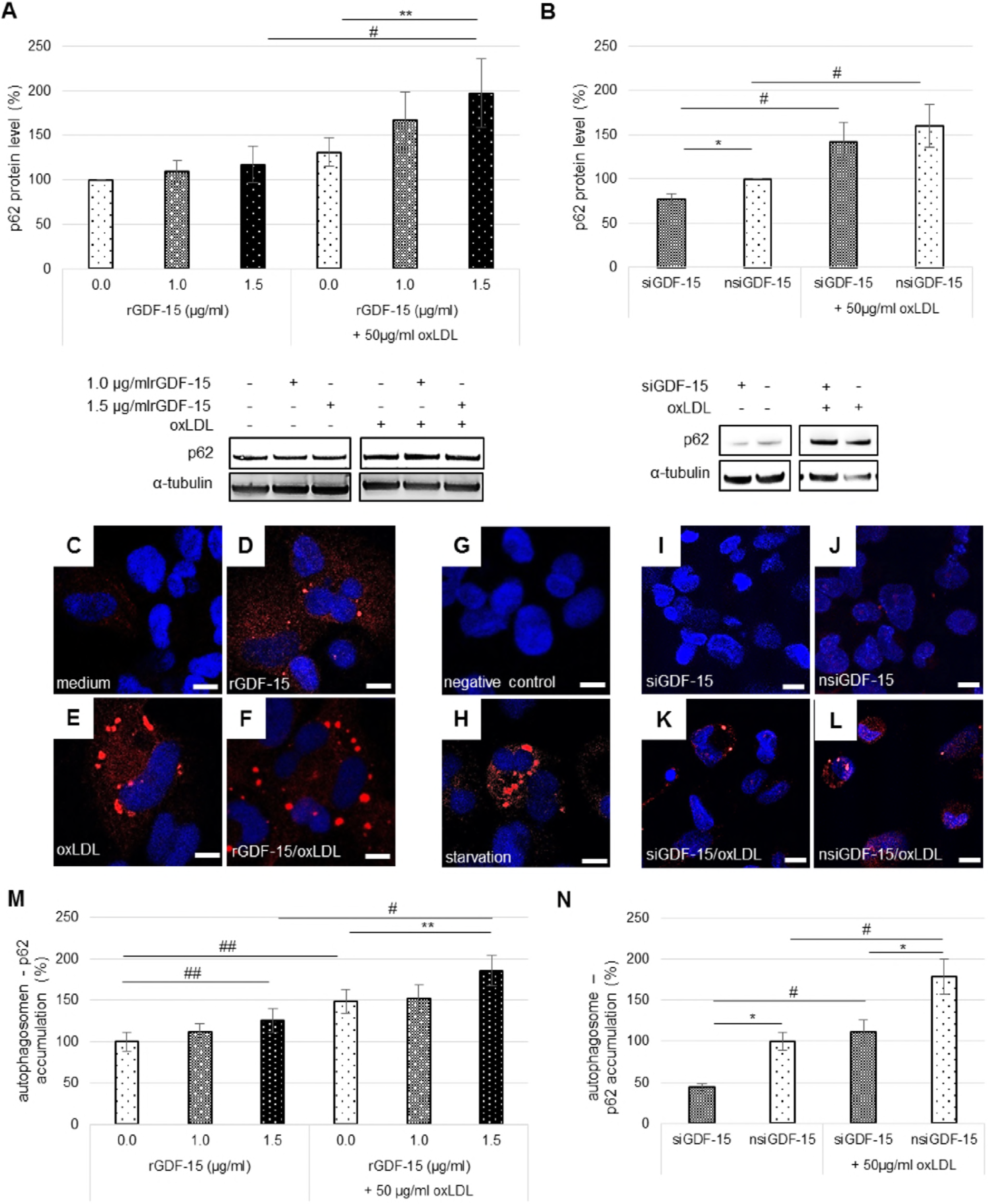
GDF-15 influences the expression and accumulation of p62 in THP-1 MΦ. THP-1 MΦ were treated with oxLDL (50 μg/ml), rGDF-15 (1.0 μg/ml, 1.5 μg/ml), oxLDL/rGDF-15 or without (medium control) (A,C-F, M). In addition siGDF-15 or nsiGDF-15 THP-1 MΦ were incubated without (medium control) or with 50 μg/ml oxLDL (B, I-L, N). A, B The p62 and α-tubulin protein levels (%) were determined using western blot analysis and quantified by ImageJ. Expression was normalized against α-tubulin (n=6-9 independent experiments). C-L Immunofluorescence of p62 accumulation in PMA-differentiated human THP-1 MΦ were observed by confocal laser scanning microscopy (Nikon Eclipse). p62 accumulation-positive control was represent by starvation (Medium+2% FKS, 4h). Negative control shows DAPI without secondary antibody against p62. The individual channels are depicted in columns: blue, DNA; red, p62. Scale bars: 10μm. M, N The p62 accumulation (%) was analyzed with Fiji ImageJ and normalized against DAPI (n=6-7 independent experiments). Data information: (A,B,M,N) Bars represent mean ± SEM. *p<0.05 significance vs. nsi GDF-15, ^#^p<0.05 significance vs. without oxLDL, **p<0.05 significance vs. oxLDL-treated MΦ, ^##^p<0.05 significance vs. medium control (Student-Newman-Keuls Method).

## Discussion

Autophagy is directly involved in lipid homeostasis (Singh ***et al***, 2009) and has been spotlighted in possible treatment of diseases associated with dyslipidemia, diabetes or atherosclerosis. GDF-15 levels are independently positively correlated with e.g. diabetes, smoking, renal dysfunction and markers of inflammation (Kim ***et al***, 2016). The cytokine GDF-15 has been hypothesized to also play a crucial role in cardiovascular diseases (Brown ***et al***, 2002; Lin ***et al***, 2014; Schlittenhardt ***et al***, 2004; Zügel ***et al***, 2010) and in the development of atherosclerosis (Bonaterra ***et al***, 2012, Johnen ***et al***, 2012; Preusch ***et al***, 2013). Moreover, clinical investigations indicate GDF-15 as a parameter for risk stratification in myocardial infarction and heart failure (Kempf ***et al***, 2006; Kempf ***et al***, 2007; Wollert ***et al***, 2007). Most recently it has been postulated that the relationship between levels of GDF-15 and inflammatory markers in atherosclerotic lesions may be more significant than that in circulation (Chen ***et al***, 2018). In relation to this, it has been shown that lifetime lack of GDF-15 considerably inhibits lumen stenosis due to less inflammation in the plaques (Bonaterra ***et al***, 2012). On the other side, it has been suggested that atherosclerotic lesions produce GDF-15 in an attempt to limit or repair the lesions and thus, when being overexpressed, GDF-15 has an overall protective effect on the atherosclerotic process (Johnen ***et al***, 2012). Therefore, our present study, which has focused on the intracellular function of GDF-15, is the first one to assign GDF-15 a pivotal role in autophagy and lipid homeostasis in human MΦ, which plays a central role during development and progression of atherosclerotic lesions.

Our study demonstrates that incubation (4h) of human THP-1 MΦ with 50 μg/ml oxLDL increased the GDF-15 mRNA expression. These data confirm earlier findings showing or suggesting a link between oxLDL-induced GDF-15 expression in human MΦ *in vitro* as well as in arteriosclerotic human carotid arteries by a co-localization of oxLDL- and GDF-15-immunoreactivity and a significant positive correlation between the percentages of oxLDL- and GDF-15-immunoreactive MΦ (Schlittenhardt ***et al***, 2004). However, we show here that rGDF-15 alone increased the cellular GDF-15 protein level in human THP-1 MΦ independent from the addition of oxLDL. Besides this, we observed that rGDF-15 – like oxLDL-increased the intracellular lipid content, independent from the addition of oxLDL. Vice versa silencing of GDF-15 in human THP-1 MΦ (with or without oxLDL) led to reduced cellular lipid content. These data clearly indicate that GDF-15 itself regulates its own expression - eventually via an autoloop feedback mechanism - and is involved in the regulation of the intracellular lipid content and consequently to foam cell formation. MΦ foam cell formation plays a central pathological role for development / progression of atherosclerotic lesions. Foam cell formation is characterized by an imbalance of cholesterol influx and efflux, accumulation of cholesterol esters (CE) within cytoplasmic lipid droplets and autophagy modulation (He ***et al***, 2017). In this context, it has been shown that the aggregation of oxLDL inhibits autophagosome elongation and sequestration (Zhou ***et al***, 2016) and induces defective MΦ autophagy (Liu ***et al***, 2016; Zhou ***et al***, 2016). Here we show that GDF-15 is able to increase the lipid content in THP-1 MΦ namely independent of oxLDL, possibly due to enhanced p62-accumulation. In line, a recent study has shown that autophagy activator rapamycin decreased intracellular lipid content during the process of foam cell formation in THP-1 MΦ markedly (Liu ***et al***, 2016). However, the role of GDF-15 during autophagic activity or autophagosome formation in THP-1 MΦ is not known to date.

Interestingly, we found that the incubation of human THP-1 MΦ with both, oxLDL plus rGDF-15 results in an increased protein level of ATG5, ATG12/ATG5 and p62, whereas silencing of GDF-15 leads to a reduction of ATG5, ATG12/ATG5 and p62 protein levels. In general, p62-protein, which regulates cell survival over packing and delivery of polyubiquitinated, misfolded, aggregated proteins and dysfunctional organelles (Pankiv ***et al***, 2007; Kim et al. 2008; Kirkin ***et al***, 2009), is used as marker of the autophagic status (Bjorkoy ***et al***, 2005; Komatsu ***et al***, 2007; Klionsky ***et al***, 2011; Mathew ***et al***, 2009). Due to the fact, that p62 protein level was increased in human THP-1 MΦ (exclusively) after co-incubation with oxLDL plus rGDF-15, this finding is interpreted as defective autophagy. It seems that rGDF-15 promotes the p62 accumulation induced by oxLDL and consequently by increasing autophagosome formation; i.e. increases of ATG5 protein level or ATG12/ATG5-complex. For these reasons the increased autophagosome formation triggered by rGDF-15 in oxLDL-treated THP-1 MΦ, increased the p62 accumulation. On the other side, we show that GDF-15 silencing in human THP-1 MΦ resulted in a decreased protein level of ATG5, ATG12/ATG5-complex and, as well as p62, including a decreased p62 accumulation, indicating that GDF-15 is involved in the regulation of these autophagy-relevant proteins. In comparison to oxLDL-treated nsiGDF-15 THP-1 MΦ, oxLDL-treated siGDF-15 THP-1 MΦ showed a reduced p62-accumulation possibly due to the decreased ATG5 level, because ATG5 and ATG12 promote autophagosome formation and are required for the induction of autophagy (Romanov ***et al***, 2012). Furthermore, it is a fact that mice with macrophages-specific deletion of autophagy-relevant Atg5, developed plaques with increased apoptosis and oxidative stress and exhibited enhanced plaque necrosis (Liao ***et al***, 2012).

The present data show that GDF-15 plays an important role during cytoprotective autophagy. Cytoprotective autophagy would constitute an initial barrier against apoptosis when stress, e.g. oxidative stress, intensity is low (Marino ***et al***, 2014). As stress conditions increase, the induction of apoptosis results in the damage of cytoprotective mechanisms by cleavage of essential ATG proteins (Marino ***et al***, 2014) and consequently an inhibition of the autophagic flux. In addition, autophagy is associated with cell fate in process of macrophages-derived foam cell formation (Liu ***et al***, 2016). Therefore, this study shows that GDF-15 is able to regulate autophagy, which may have an important pathophysiological consequence in atherosclerosis.

## Material and Methods

### LDL-oxidation

Native (n)LDL (RayBiotech, GA, USA) oxidation was performed as previously described by Galle and Wanner (Galle & Wanner, 1998) and Steinbrecher (Steinbrecher, 1987). The nLDL was suspended in endotoxin-free phosphate-buffered saline (PBS) without Ca^2+^, Mg^2+^ (LONZA, Ratingen, Germany) to a final concentration of 1 mg protein/ml and dialyzed using Slide-A-Lyzer Dialysis Cassettes 7K MWCO (Thermo Fisher Scientific Inc., Rockford, USA). OxLDL was obtained by oxidizing nLDL using CuSO_4_ (50 μM in PBS Ca^2+^, Mg^2+^ free, 24 h, RT). Different methods verified the grade of oxidation: (1) - Trinitrobenzene sulfonic acid (TNBSA, Thermo Fisher Scientific Inc., Rockford, USA), which measures free amino groups (Foxx ***et al***, 2008). (2) - Relative electrophoretic mobility (REM) by agarose gel electrophoresis and visualized by staining with Coomassie Blue (Lougheed & Steinbrecher, 1996), and (3) - by spectrophotometric analysis (absorbance spectrum between 400 and 700nm) (Galle & Wanner, 1998). OxLDL was used by an increased REM of 17.5 % ± 3.34 % compared to nLDL, an increased percentage of blocked amino groups (40.4 % ± 0.65 % compared to LDL) and by the disappearance of the nLDL characteristic absorption peaks at 460 and 485 nm (Galle & Wanner, 1998) (Figure EV1).

### Cell culture, transfection and gene silencing

The human leukemic monocyte cell line THP-1 (Leibniz Institute DSMZ, Braunschweig, Germany) was used. Originally, the culture was derived from the blood of a one-year old boy with acute monocytic leukemia and is frequently used as a model of monocyte/MΦ cell lineage (Tsuchiya ***et al***, 1980). THP-1 cells were cultured in RPMI-1640 medium (Capricorn Scientific GmbH, Ebsdorfergrund, Germany) supplemented with penicillin and streptomycin (Capricorn Scientific GmbH,) and 10% fetal bovine serum (Capricorn Scientific GmbH,). Cells were cultured at 37°C in a 5% C0_2_ environment with a medium change every 2-3 days. All experiments were performed using cells at passage 9 and lower. RPMI 1640 medium supplemented with 100 nM Phorbol 12-mystriate 13-acetate [PMA, (Sigma-Aldrich Chemie GmbH Munich, Germany)] was used (72h) to THP-1 cells to induce monocyte differentiation into MΦ. Transfection of THP-1 MΦ with 50 nM small interfering RNA (siRNA) for GDF-15 (FlexiTube GeneSolution GS9518, QIAGEN GmbH, Hilden, Germany) and with negative siRNA (nsiGDF-15) (AllStars Negative Control, QIAGEN GmbH) was performed using HiPerfect Transfection Reagent (QIAGEN GmbH) following the manufacturer’s instructions. As positive control, we used the AllStars Hs Cell Death Control siRNA (QIAGEN GmbH), a compound of highly potent siRNAs targeting ubiquitously expressed human genes that are essential for cell survival. Transfection efficiency was estimated by observing cells by light microscopy 48 h after transfection with the AllStars Hs Cell Death Control siRNA. After transfection, cells were treated with 50 μg/ml oxLDL to induce foam cell formation or left untreated (medium) for 4h. PMA-differentiated THP-1 MΦ were treated with 1.0 - 1.5 μg/ml human rGDF-15 [rGDF-15 (ProVitro AG, Berlin, Germany)] or co-incubated with oxLDL+rGDF-15 for 4h.

### GDF-15 ELISA

For the quantification of human GDF-15 we used the DuoSet® ELISA Development System (R&D Systems, Inc., Abingdon, UK). The Capture Antibody was coated at a 96-well MaxiSorp-ELISA Microplate (Nunc, San Diego, USA) and incubated overnight at room temperature. According to manufacturers’ instructions, after the blocking step, the samples (2.5 μg protein/well) or standards were added into the well. *After the incubation of a detection antibody and Streptavidin-HRP, we added the substrate solution (SigmaFast*™ *OPD, Sigma-Aldrich Chemie GmbH) to each well and incubated for 30 minutes in the dark. After that, the reaction was stopped with 50 μl 3 M HCl. The GDF-15 protein level (pg/ml) was measured by an ELISA reader (Tecan Deutschland GmbH, Crailsheim, Germany) at OD_490/655 nm_*.

### SDS-PAGE and Western blot

After the treatments, PMA-differentiated THP-1 MΦ were washed in ice-cold PBS and lysed using radioimmuniprecipitation assay (RIPA) buffer pH 7.5 (Cell Signaling Technology, Frankfurt, Germany), containing protease/phosphatase inhibitor cocktail (Cell Signaling Technology). The protein concentrations were determined spectrophotometrically using the Pierce BCA (bicinchoninic acid) Protein Assay (Thermo Scientific, Rockford, USA). Proteins were loaded on NuPAGE® Novex® 412% Bis-Tris Gels, pre-cast polyacrylamide gels (Life Technologies GmbH, Darmstadt Germany). Proteins were transferred onto 0.45 μm nitrocellulose membranes (Millipore, Billerica, MA, USA). Primary Antibodies (Appendix Table S1) were added and incubated overnight at 4°C in blocking buffer (5% fat-free milk). Membranes were incubated with enhanced ECL-anti-goat IgG-POD antibody, ECL-anti-mouse IgG-POD antibody or ECL-anti-rabbit IgG-POD antibody. The peroxidase reaction was visualized by AceGlow chemiluminescence substrate (PEQLAB GmbH, Erlangen, Germany) and documented by the Fusion-SL Advance™ imaging system (PEQLAB GmbH) according to the manual instructions. The intensity of the specific western blot bands was quantified using the software ImageJ from the National Institutes of Health (Bethesda, USA) and normalized against α-tubulin.

### Reverse transcription and quantitative polymerase chain reaction

Total RNA was extracted from human THP-1 MΦ using PeqGold TRIFast™ (PEQLAB GmbH). DNase I (RNase-free; Thermo Scientific) was used according to the manufacturer′s instructions. The AffinityScript Multiple Temperature Reverse Transcriptase and the Brilliant III Ultra-Fast SYBR® Green Master Mix were obtained from Agilent Technologies Deutschland GmbH. (Waldbronn, Germany). All primers were purchased from QIAGEN GmbH (Hilden, Germany) (Appendix Table S2). RNA concentration and purity were determined by absorbance measurements at 260 nm and 280 nm (A260/A280=1.7-2.0) using a NanoDrop 8000 Spectrophotometer (Thermo Scientific, Schwerte, Germany). Total RNA integrity was confirmed by lab-on-a-chip technology, using an RNA 6000 NanoChip kit on an Agilent 2100 Bioanalyzer (Agilent Technologies Deutschland GmbH). RNA was only used with a RNA Integrity Number (RIN) of 8.5 ± 0.9. 1.0 μg of template RNA was used for cDNA synthesis; RNA was reverse transcribed using Oligo(dT)_12-18_ primer and the AffinityScript™ Multiple Temperature Reverse Transcriptase, according to the manufacturer’s instructions. The cDNA (diluted 1:20) was amplified using the Brilliant III Ultra-Fast SYBR® Green QRT-PCR Master Mix (Stratagene-Agilent Technologies, Waldbronn, Germany). Amplification and data analyses were performed using the M×3005P™ QPCR System (Stratagene). The data were analyzed using the relative standard curve method. The NormFinder software program was used to ascertain the most suitable reference gene to normalize the RNA input as described earlier (Stern-Streater ***et al***, 2009).

### Oil Red O staining

The Oil Red O [ORO, (Sigma-Aldrich Chemie GmbH)] working solution was prepared by diluting the stock solution (3 mg/ml ORO dissolved in 2-propanol) with distilled water (3:2). For staining, PMA differentiated THP-1 MΦ were fixed in 10% paraformaldehyde (PFA), in PBS and after washing with PBS the ORO working solution was added to the culture dishes afterwards. The nuclei were counterstained using Crystal violet (Carl Roth GmbH, Karlsruhe, Germany).

### Immunocytofluorescence confocal laser scanning microscopy

Cells were fixed with ice-cold methanol and permeabilized with 0.1% Triton-X 100 in PBS. Thereafter, the detergent was removed by repeated washing in PBS. Primary antibodies (Appendix Table S1) were applied in PBS overnight (4°C). After incubation with secondary antibodies (Appendix Table S1) and subsequent staining with DAPI, we covered the cells with Immu-Mount™ (Thermo Electron Corporation; Pittsburgh; USA). Images were taken by confocal laser-scanning-microscope Eclipse Ti-E, (Nikon GmbH, Düsseldorf, Germany).

### Statistical analyses

Statistical analyses were performed using SigmaPlot 12 (Systat Software Inc., USA). After testing for normality (by Shapiro-Wilk), the unpaired Student’s t-test or one-way analysis of variance (ANOVA) was used. Data are reported as mean ± standard error of the mean (SEM). p<0.05 was considered as statistically significant.

## Acknowledgement

We thank Anne Henkeler, Andrea Cordes, Elke Völck-Badouin and Gabriella Stauch for excellent technical assistance.

## Author contribution

Conception and design of the study: AS, KR. Acquisition and analysis of data: KA, AS. Drafting the manuscript: KA, AS. Critical review of manuscript: RK, GB.

## Expanded View

Figure S1 - Modification of nLDL with 50MM CuSO4 results to mid-oxidizied LDL.

(A) The relativ elektromobility shift (REM) of BSA and mid-oxidizied LDL in relation to nLDL.

(B) Blocking of amino groups was calculated by comparing the free amino groups remaining in oxLDL to those of the nLDL. The percentage of blocked amino groups of mid-oxidizied LDL compared to nLDL.

(C) Represenative spectrophotometric analysis of nLDL and mid-oxidizied LDL using wave length 400 – 700 nm.

Data information: (A) Bars represent means ± SEM of 7 independent experiments (B) Bars represent means ± SEM of 3 independent experiments (A, B)*p<0,001 vs. nLDL (OneWayANOVA, Holm-Sidak-method).

Table S1 - Antibodies used in this study.

Table S2 - Primers used in this study.

